# Efficient production of Moloney murine leukemia virus-like particles pseudotyped with the severe acute respiratory syndrome coronavirus-2 (SARS-CoV-2) spike protein

**DOI:** 10.1101/2020.09.16.298992

**Authors:** Sylvie Roy, Karim Ghani, Pedro O. de Campos-Lima, Manuel Caruso

**Affiliations:** CHU de Québec-Université Laval Research Center (Oncology division), Université Laval Cancer Research Center, Québec, Qc, Canada; BioVec Pharma, Québec, Qc Canada; Boldrini Children’s Center, Campinas, Brazil; Functional and Molecular Biology Graduate Program, State University of Campinas, Brazil; Department of Molecular Biology, Medical Biochemistry and Pathology, Faculty of Medicine, Université Laval, Québec, Qc, Canada

## Abstract

The severe acute respiratory syndrome coronavirus 2 (SARS-CoV-2) outbreak that started in China at the end of 2019 has rapidly spread to become pandemic. Several investigational vaccines that have already been tested in animals and humans were able to induce neutralizing antibodies against the SARS-CoV-2 spike (S) protein, however protection and long-term efficacy in humans remain to be demonstrated.

We have investigated if a virus-like particle (VLP) derived from Moloney murine leukemia virus (MLV) could be engineered to become a candidate SARS-CoV-2 vaccine amenable to mass production. First, we showed that a codon optimized version of the S protein could migrate efficiently to the cell membrane. However, efficient production of infectious viral particles was only achieved with stable expression of a shorter version of S in its C-terminal domain (ΔS) in 293 cells that express MLV Gag-Pol (293GP). The incorporation of ΔS was 15-times more efficient into VLPs as compared to the full-length version, and that was not due to steric interference between the S cytoplasmic tail and the MLV capsid. Indeed, a similar result was also observed with extracellular vesicles released from parental 293 and 293GP cells. The amount of ΔS incorporated into VLPs released from producer cells was robust, with an estimated 1.25 *μ*g/ml S2 equivalent (S is comprised of S1 and S2). Thus, a scalable platform that has the potential for production of pan-coronavirus VLP vaccines has been established. The resulting nanoparticles could potentially be used alone or as a boost for other immunization strategies for COVID-19.

**IMPORTANCE:** Several candidate COVID-19 vaccines have already been tested in humans, but their protective effect and long-term efficacy are uncertain. Therefore, it is necessary to continue developing new vaccine strategies that could be more potent and/or that would be easier to manufacture in large-scale. Virus-like particle (VLP) vaccines are considered highly immunogenic and have been successfully developed for human papilloma virus as well as hepatitis and influenza viruses. In this study, we report the generation of a robust Moloney murine leukemia virus platform that produces VLPs containing the spike of SARS-CoV-2. This vaccine platform that is compatible with lyophilization could simplify storage and distribution logistics immensely.

---

A cluster of severe pneumonia cases emerged in Wuhan in the Chinese province of Hubei in December 2019 and has quickly become a worldwide pandemic. A new virus was later identified as the etiological agent: the severe acute respiratory syndrome coronavirus 2 (SARS-CoV-2), and the condition was named coronavirus disease 2019 (COVID-19) by the World Health Organization (1, 2). As of today, September 16^th^ 2020, 29 million people have been infected, and 939,000 deaths have been recorded (gisaid.org), but these numbers are probably well underestimated. In addition to its severe health threat, COVID-19 has profound socioeconomic consequences (3).

SARS-CoV-2 is the seventh coronavirus that has been identified so far. HCoV-NL63, HCoV-229E, HCoV-OC43 and HKU1 strains are constantly present in the human population and cause mild common-cold symptoms (4). The other two, SARS-CoV and the Middle East respiratory syndrome (MERS)-CoV, are similar to SARS-CoV2 in that they are highly pathogenic to humans causing acute respiratory disease (5). Two epidemics were caused by SARS-CoV and MERS-CoV, respectively: SARS that originated in China in 2002 and MERS which emerged 10 years later in the Middle East. These two viruses did not spread widely as only 8,096 cases were reported for SARS-CoV and 2,494 for MERS but had an exceedingly high mortality rate (9-35%). There have been no new cases of SARS-CoV reported since 2004 although MERS is still endemic in the Middle East (6). The three highly pathogenic coronaviruses are zoonotic and have emerged from bats with dromedary camels, palm civet and most likely pangolin being the intermediary host for MERS-CoV, SARS-CoV and SARS-CoV-2, respectively (4, 7–13). Coronaviruses are single-stranded positive-sense RNA viruses that are composed of four structural proteins: spike (S), nucleocapsid, envelope and membrane (4). The S protein that is about 180 kDa assembles as a trimer at the virus surface. It is composed of two subunits S1 and S2 that are responsible for the virus attachment and fusion. MERS uses dipeptidyl peptidase 4 as its receptor, while SARS-CoV and SARS-CoV-2 share the same receptor for entering cells: the angiotensin-converting enzyme 2 (ACE2) (13–19).

Efforts are being made to identify candidate neutralizing antibodies (Nabs) that could block the interaction of SARS-CoV2-S with its receptor and that could be used for treating infected patients (20–22). Several vaccine strategies for COVID-19 are also intensively pursued, with S protein being the major target (23–25). These vaccines are produced from different platforms: RNA, DNA, recombinant proteins, viral vector-based or virus-like particles (VLPs), and live attenuated and inactivated viruses (23–25).

Vaccines made from RNA, DNA or proteins are usually easier to manufacture than those that are virus-derived but it is generally accepted that vaccines made of the original virus (attenuated) or from VLPs induce a better immune response (26). This is an important point to consider as a COVID-19 vaccine ideally should induce high-titer Nabs for a long-lasting period of time.

Preliminary results obtained in animals and in humans have shown that both humoral and cellular immune responses can be obtained with different vaccine strategies, and that Nab titers achieved by vaccination in humans were comparable to those measured in the serum of COVID-19 convalescent individuals (25, 27–38). A recent study evaluating a DNA vaccine indicated that macaques were protected upon SARS-CoV-2 challenges 13 weeks after vaccination (39). However, only long-term studies in humans will tell us about the efficacy of all these vaccines.

VLPs are produced by the assembly of viral proteins that do not contain genetic material, and that are then unable to replicate. VLPs are advantageous for their immunostimulatory activity: they are highly recognized by antigen-presenting cells and the repetitive arrangement of antigens on their surface is capable of inducing both innate and adaptive immune responses with a high level of Nabs (26). VLPs have already been successfully developed for Human Papilloma Virus, Hepatitis B, E, and A Virus and influenza virus (26).

The difficulty of developing COVID-19 vaccines in a short period of time is compounded by the major hurdle of creating mass production capacity to deliver the final product for the entire world population. In this study, we have engineered and characterized a Moloney murine leukemia virus (MLV) VLP platform that has the potential for large-scale production of a COVID-19 vaccine.

## RESULTS

### The SARS CoV-2 S protein migrates to the cell surface

The production of an MLV-derived VLP COVID-19 vaccine requires the presence of the carried immunogenic molecule (in our case the S protein) at the surface of the producer cell. As coronaviruses assemble at the ER-Golgi intermediate compartment (4), we had first to verify if the S protein could migrate efficiently to the cell surface. A full-length, codon-optimized S gene as well as a shorter version in which the last 3’ 57 nucleotides are lacking were cloned into an expression vector. The rationale for the construction of the latter is based on the presence of an endoplasmic reticulum retention signal in the cytoplasmic tail of the coronaviruses S protein and previous reports that a 19-amino acid C-terminal deletion of SARS-CoV S increases the production of MLV or vesicular stomatitis (VSV) infectious particles (40–46). After transfection in 293 cells, both S versions were detected with a very similar intensity at the cell surface (Fig. 1). These results indicated that S was able to efficiently migrate to the cell membrane and that, in these experimental conditions, the endoplasmic reticulum retention signal did not affect its localization.

**FIG 1.**
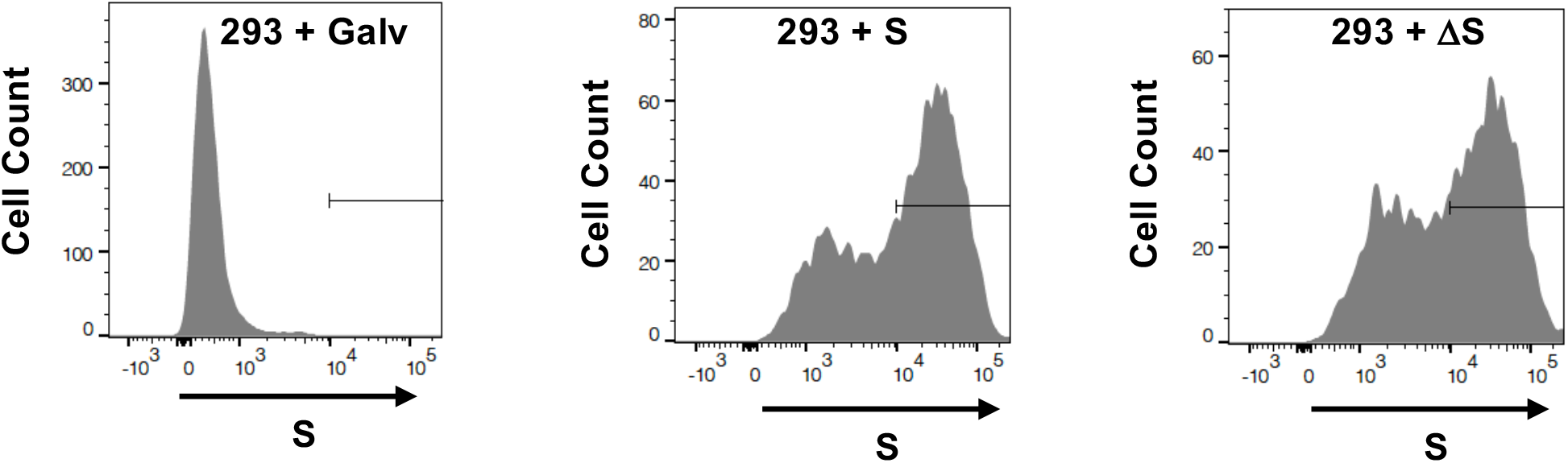
Expression of S protein at the surface of 293 cells. FACS analysis of cells transiently transfected with plasmids encoding the Galv envelope, the full-length S protein, and the ΔS version. S was detected with an anti-S1 antibody.

### Inefficient transient production of infectious recombinant MLV viruses pseudotyped with the SARS CoV-2 S protein

The production of VLPs pseudotyped with S or ΔS (VLP-S) was next assessed by generating GFP recombinant viruses in transient transfections. Titers were measured by FACS analysis on 293-ACE2 cells, a cell line generated by stable transfection that is 61% positive for ACE2 (Fig. 2). Titers of 3.2 × 10^7^ infectious units (IU)/ml and 1.5 × 10^6^ IU/ml were obtained for VSV-G- and Galv-pseudotyped viruses, although titers of S and ΔS-pseudotyped viruses were below the detection limit of 10^4^ IU/ml (Fig. 3A). Only few GFP cells could be observed by fluorescence microscopy after infection with the ΔS-pseudotyped virus and there were none when the S-pseudotyped vector was used (Fig. 3B). Thus, these results indicated that the transient production was extremely inefficient for generating VLP-S, even with ΔS.

**FIG 2.**
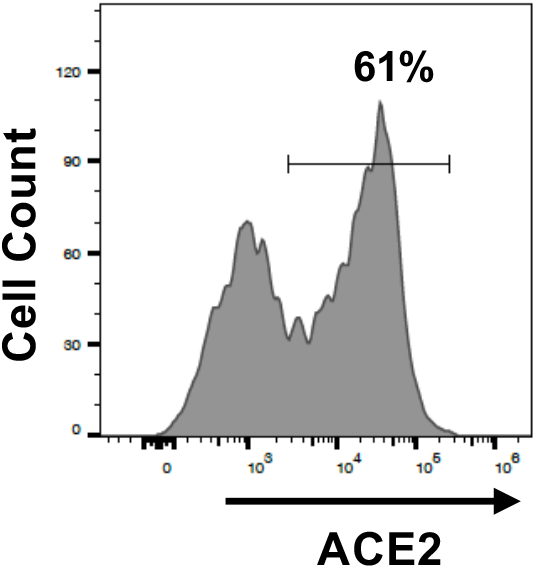
Expression of ACE2 at the surface of 293-ACE2 cells measured by FACS analysis.

**FIG 3.**
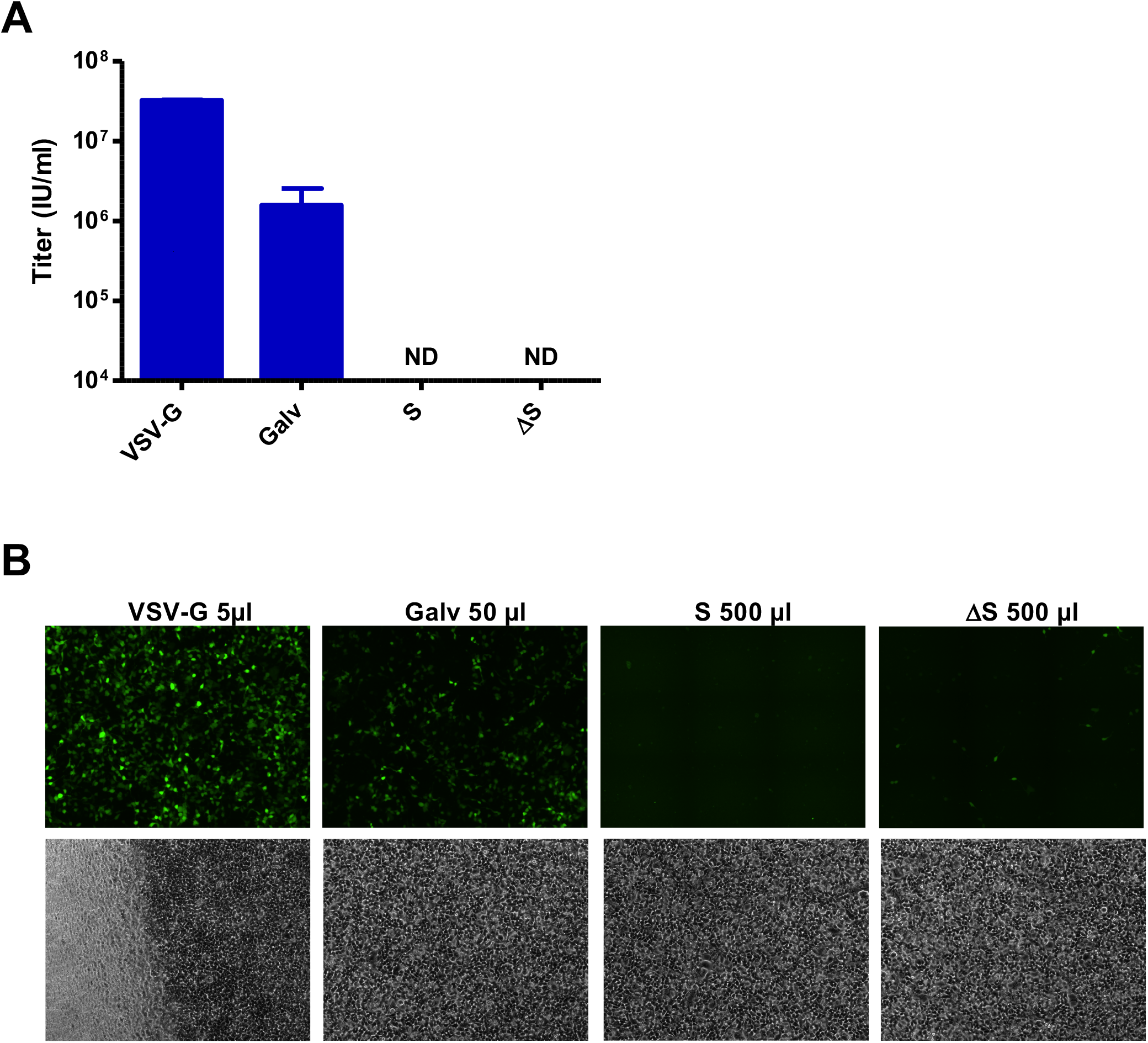
Transduction efficiency of different GFP pseudotyped vectors produced in transient transfections. Two days after infection of 293-ACE2 cells, titers of VSV-G-, Galv-, S- and ΔS-pseudotyped vectors were (A) measured by FACS analysis or (B) evaluated by fluorescence microscopy. Values presented are the mean ± SD of three independent experiments. Fluorescent and bright-field pictures are displayed. The envelope pseudotype and the volume used for infection are indicated.

### ΔS-pseudotyped MLV recombinant viral particles are efficiently released from stable producer cells

We have shown that stable retrovirus packaging cell lines can generate Galv-pseudotyped vectors with at least 10-fold higher titers as compared to transient transfection productions (47). We then hypothesized that S or ΔS stably expressed in 293GP cells (293 cells that express MLV Gag-Pol) could be a better system to produce VLP-S. Stable populations of 293GP cells expressing S and ΔS were then generated by transfection. In these cells, S and ΔS were able to localize at the cell surface at even higher levels than what we found in transient transfection (Fig. 4A). A GFP retroviral vector was then introduced in these cells by infection (Fig 4B), and titers of GFP viruses released by these new producers were measured after infecting 293-ACE2 cells. Only few GFP positive cells could be detected by fluorescence microscopy after infection of 293-ACE2 cells with the S-pseudotyped vector, but a very high percentage of fluorescent cells was observed after infection with the ΔS virus. A high number of GFP positive cells was seen with the Galv virus diluted 10-times as compared to the two other vectors (Fig. 5A). Titers of 1.6 × 10^7^ IU/ml and 10^5^ IU/ml were measured for the Galv and ΔS-pseudotyped viruses, respectively, and the S-pseudotyped vector titer was below the detection limit of 10^4^ IU/ml, as expected (Fig 5B). We could conclude that the production of recombinant viral particles was robust from stable producers expressing ΔS and inefficient with the full-length version of SARS CoV-2 S.

**FIG 4.**
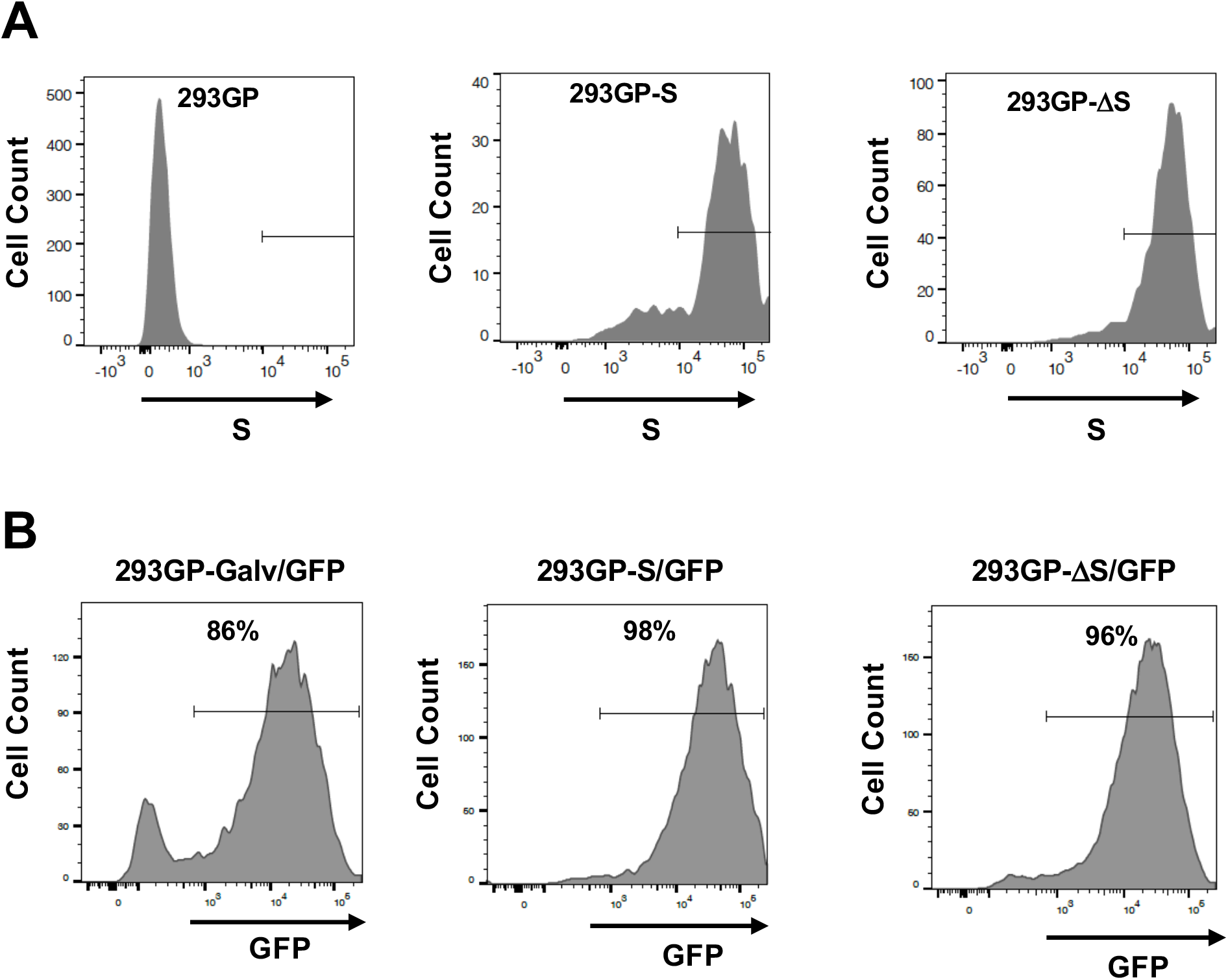
Characterization of stable VLPs producer cells. (A) S expression was measured by FACS analysis of 293GP, 293GP-S and 293GP-ΔS cells with an anti-S1 antibody. (B) GFP fluorescence of 293GP-Galv/GFP, 293GP-S/GFP and 293GP-ΔS/GFP measured by FACS analysis.

**FIG 5.**
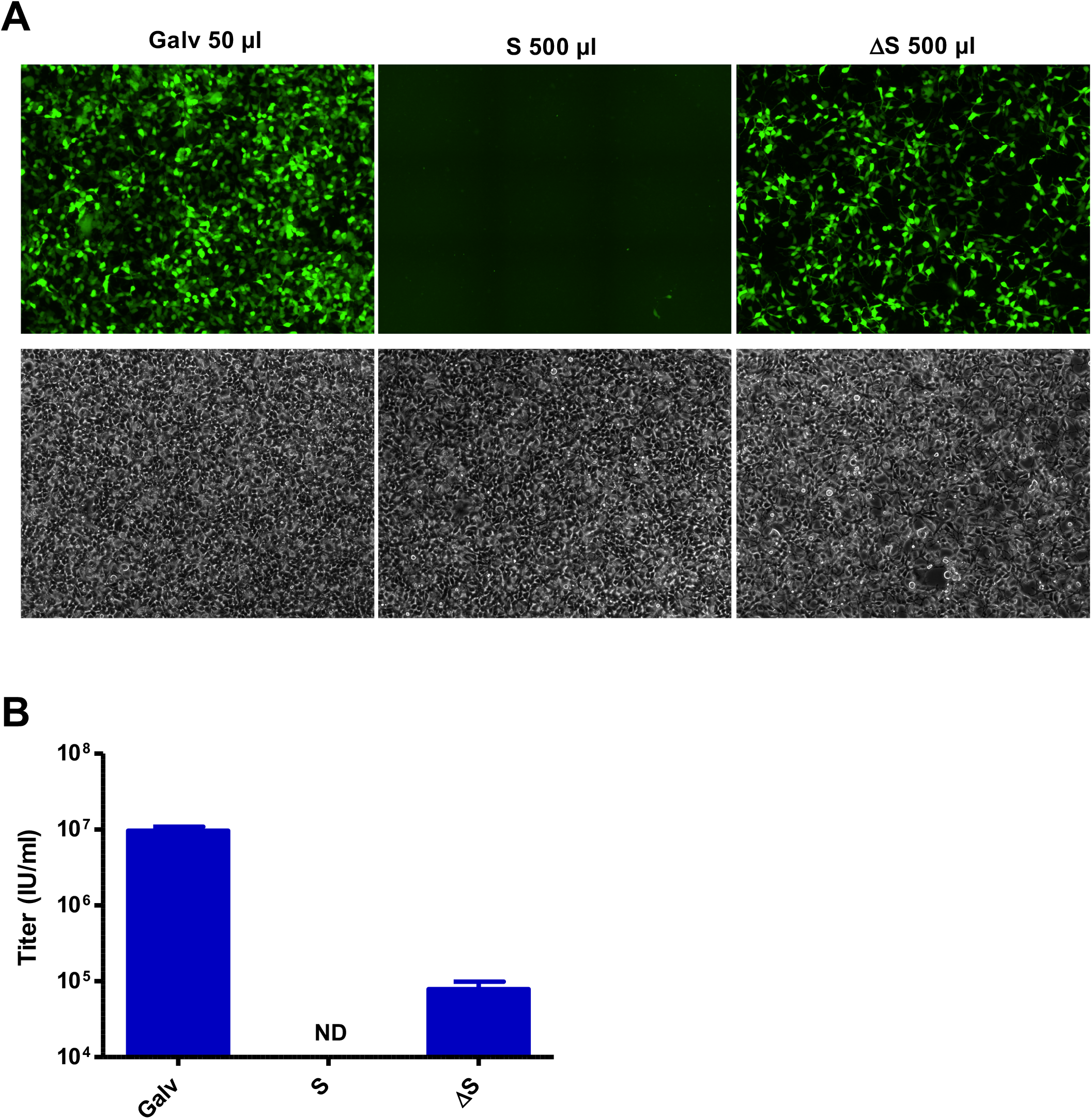
Transduction efficiency of GFP pseudotyped vectors released from stable producers. (A) Fluorescent and bright-field pictures are displayed. The envelope pseudotype and the volume used for infection are indicated. (B) Titers of Galv-, S- and ΔS-pseudotyped vectors produced from stable producers were measured by FACS analysis two days after infection. Values presented are the mean ± SD of three independent experiments.

### The deletion of the 19 amino acid cytoplasmic tail of S does not enhance its fusogenicity

As producer cells express the same amount of S and ΔS at the cell surface, one possible explanation for the high transduction efficiency of ΔS-pseudotyped vectors could be increased fusogenicity. The fusion capacity of S and ΔS was then assessed in a syncytia formation assay by mixing 293GP cells expressing S or ΔS with 293-ACE2 cells. The number and the size of syncytia evaluated one day after mixing were very similar between S and ΔS mixtures, and there were none with the control 293 cells (Fig. 6). Thus, the deletion of the 19 amino acids in the S cytoplasmic tail does not have a significant effect on its fusogenicity.

**FIG 6.**
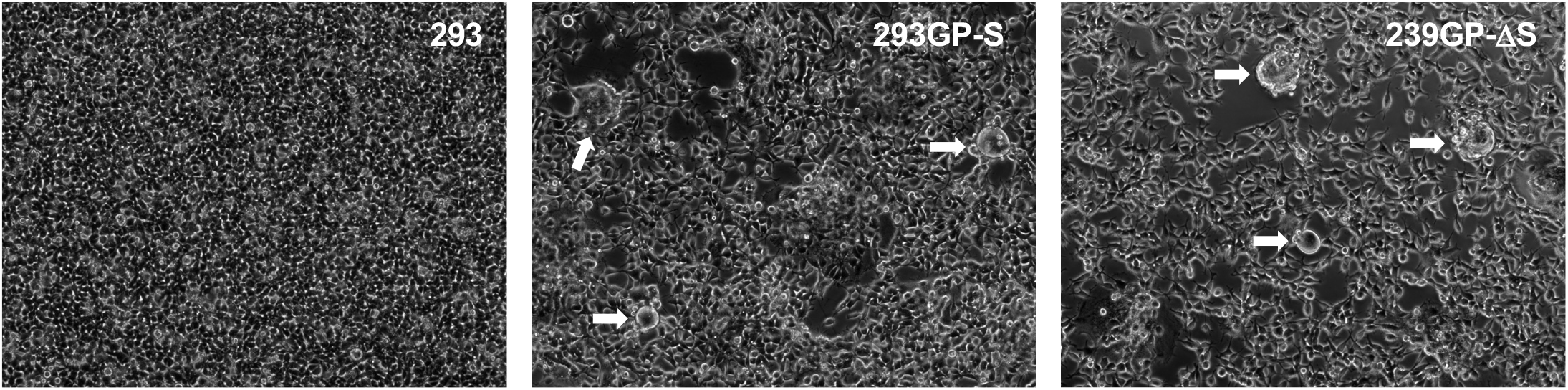
Fusion mediated by S and ΔS. 293, 293GP-S and 293GP-ΔS were mixed with 293-ACE2 cells at a 1/10 ratio. Syncytia (arrow) were observed 24 h later.

### High amounts of SARS-CoV-2 ΔS protein are incorporated into MLV VLPs released from stable producer cells

A VLP-derived SARS CoV-2 vaccine will be a viable option if sufficient amounts of S protein are incorporated at the surface of the released particles. Western blots were performed with an anti-S2 antibody to evaluate the quantity of S protein into VLPs produced in transient transfections and from stable producers. Two bands were detected around 90 KDa that are most likely two glycosylated forms of S2. The uncleaved S protein migrated around 180 kDa, and two other bands above 250 kDa were also detected in the ΔS samples that had more intense signals. These bands could be dimeric and trimeric forms of S as it has been suggested (19). The amount of S2 detected at the surface of VLPs produced in transient transfections or released from stable producers was much higher with the truncated version of S than with the full-length molecule (Fig. 7A). MLV viral particles produced in transient transfection or from stable producers were detected with an antibody against p30. A 4- and a 15-fold difference was found with the transient and the stable production systems, respectively (Fig. 7B), although there was less than a 1.5-fold difference between S and ΔS in cellular extracts (Fig. 7C). More ΔS was also released as compared to the full-length protein in the supernatants of stably transfected 293 cells, however the amount of ΔS detected was 4-to-5 times lower than the one released from the 293GP-ΔS. The amount of S2 equivalent present in the supernatant of 293GP-ΔS cells was high and evaluated at 1.25 μg/ml using the IgG-S2 standard (Fig. 7A). Our results indicated that the incorporation of S into MLV VLPs is very efficient in stable producers but only with the truncated version of S.

**FIG 7.**
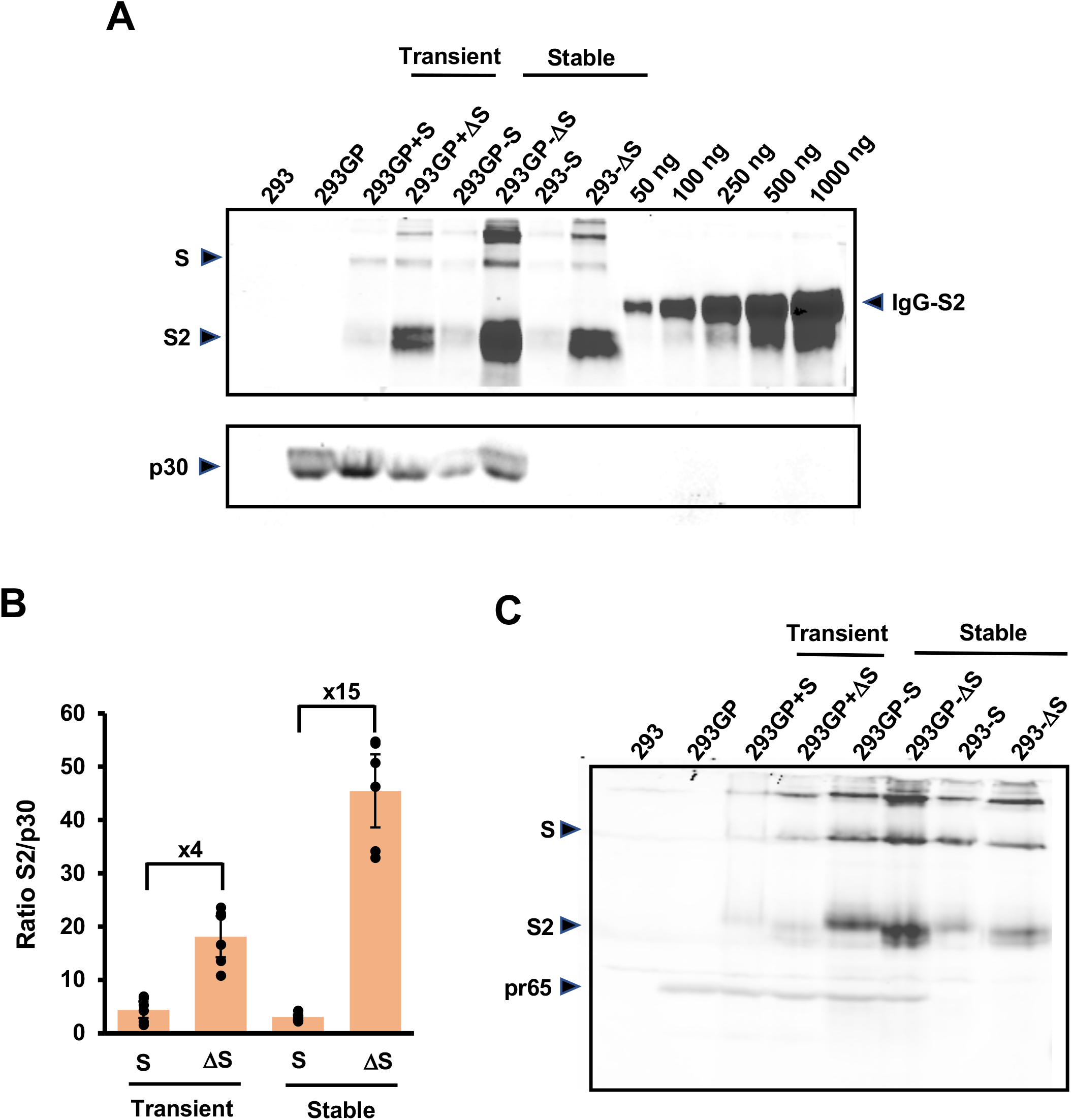
Quantification of S and ΔS incorporated into VLPs. (A) Western blot analysis from concentrated supernatants of 293GP and 293 cells using anti-S2 and anti-p30 antibodies. Different amounts of Fc-tagged S2 were also loaded on the gel to quantify S2 in VLPs. (B) Differences between S and ΔS incorporation into VLPs. All the bands detected by the anti-S2 antibody in S- and ΔS-containing samples were quantified and normalized with the signal obtained for MLV p30. Values presented are the mean ± SD of three independent experiments analyzed twice in Western blot. (C) Western blot analysis of SARS CoV-2 S protein in cellular extracts. Signals for S2, S, and multimeric forms of S were detected with the anti-S2 antibody. The Gag precursor pr65 was detected with the anti-p30 antibody.

### SARS-CoV2 ΔS protein is preferentially incorporated into MLV VLPs versus extracellular vesicles

As stable transfected 293 cells were capable of releasing S or ΔS, we decided to further characterize the supernatants of the 293GP-ΔS. We used an iodixanol velocity gradient to discriminate VLPs from extracellular vesicles (EVs) as this technique has been used in the past to successfully separate human immunodeficency viruses (HIV) from EVs (48, 49). Western blots with anti-S2 and anti-p30 antibodies were performed on the collected gradient fractions of 293-ΔS and 293GP-ΔS supernatants. S2 was detected in the top fractions from the 293-ΔS supernatant but there were none in the last 3 bottom fractions (Fig. 8A). A similar detection pattern was observed in the top fractions of the supernatant from 293GP-ΔS, but the majority of S2 came from the 2 bottom fractions in which a band corresponding to the uncleaved S protein was also detected. The p30 signal was present in these two fractions, which indicated that the majority of ΔS released from 293GP-ΔS cells was incorporated into VLPs (Fig. 8B).

**FIG 8.**
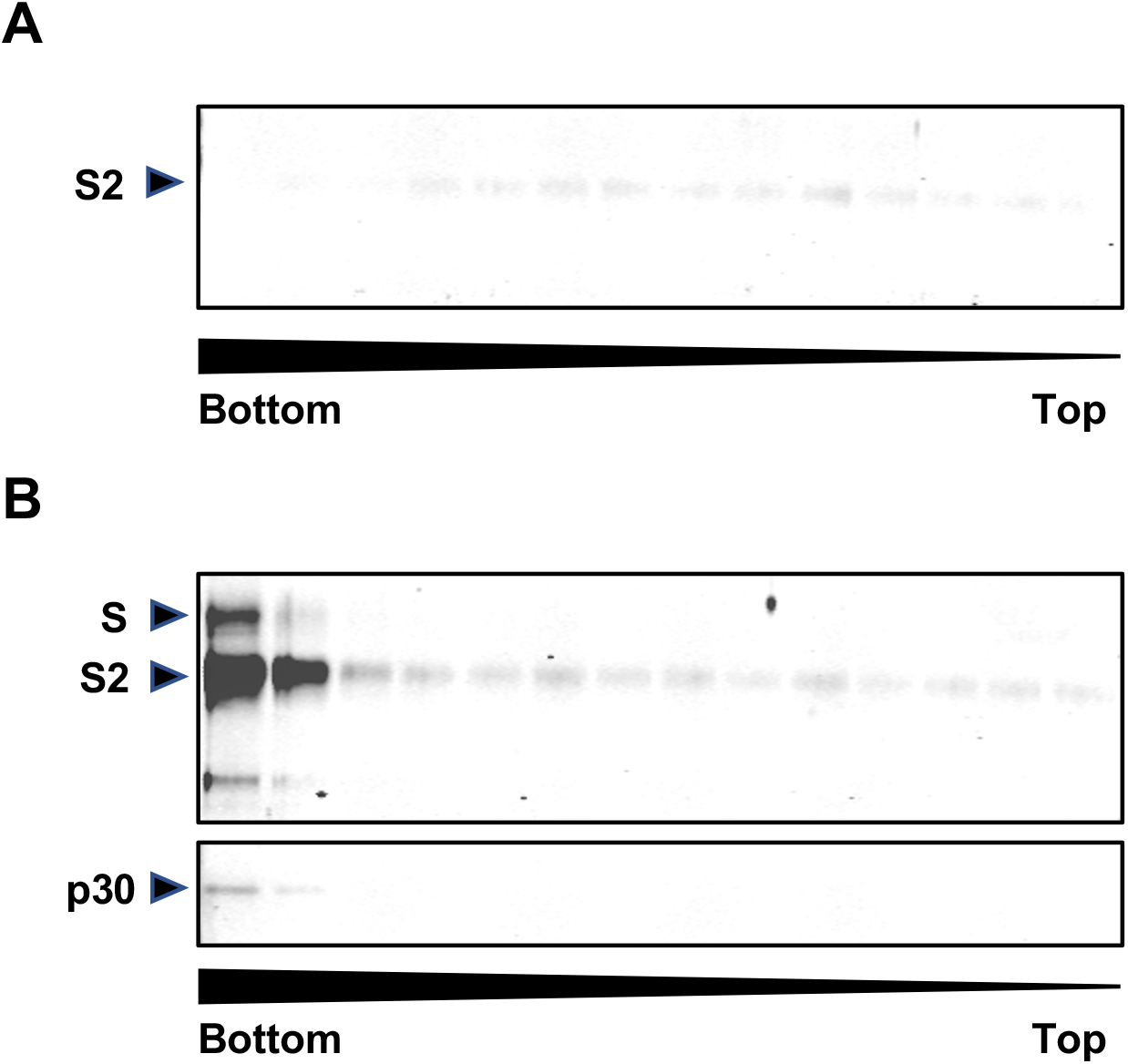
Incorporation of SARS-CoV-2 ΔS into MLV VLPs. Western blot analysis with antibodies against S2 and p30 on collected fractions separated with an iodixanol velocity gradient of (A) 293-ΔS and (B) 293GP-ΔS supernatants. The arrow below the blot indicates the density gradient.

## DISCUSSION

Immunization will be the best preventive strategy to address the current COVID-19 pandemic, although therapeutic alternatives cannot be neglected as an efficient vaccine is not a certainty (23, 25, 50). Yet preliminary results from preclinical and clinical studies are encouraging as several types of vaccines are able to trigger the production of Nabs against SARS-CoV-2 S (25, 27–31, 33–39, 51, 52). How efficient and how long these Nabs will be present in vaccinated people remains an open question that will only be answered with time (25). Also, antibody-dependent enhancement will have to be carefully monitored in these trials as it is a side effect that cannot be underestimated with coronaviruses (23, 25, 50). One other major challenge ahead will be the capacity to mass produce COVID-19 vaccines. In this study, we have established and characterized a new MLV-derived VLP platform that could be used for the production of a COVID-19 vaccine.

The efficient pseudotyping of MLV particles with S is a prerequisite to establish a robust VLP platform. Studies on SARS-CoV and more recently on SARSCoV-2 have shown that the codon optimization of S and the deletion of the ER retention signal located in the cytoplasmic tail are modifications that increase the pseudotyping of MLV, HIV, simian immunodeficiency and VSV viral vectors (40, 46, 53–55). The codon optimization enhances the level of S expression, but the role of the ER retention signal is less clear. Indeed, it was recently reported that the localization of SARS-CoV-2 S at the cell surface was not improved after disrupting the ER retention signal by missense mutations (56). In this study, we showed that S could be detected at the cell surface at a similar level to that achieved by ΔS in transiently transfected cells as well as in stable producers (Fig. 1 and Fig. 4A), a finding that has also been reported for SARS-CoV S expressed in transient transfections (40, 54). These results indicate that S can bypass its natural localization and efficiently migrates to the cell surface when it is overexpressed.

Despite similar amounts of S and ΔS at the cellular membrane, the truncated version was more efficiently incorporated into MLV viral particles. Four- and 15-fold differences were obtained with VLPs produced in transient transfection experiments and from stable producers, respectively (Fig. 7B). The hypothesis that has been proposed for SARS-COV and SARS-CoV-2 is that the 19 amino-acid deletion in the S cytoplasmic tail facilitates the pseudotyping by decreasing the steric interference with the retroviral matrix proteins (54, 55, 57). Our results invalidate this hypothesis as more ΔS was also found in the supernatant of 293 transfected cells that did not express MLV Gag-Pol (Fig. 7A). Parental 293 and 293GP cells release EVs that can incorporate ΔS more efficiently than S (Fig. 7A and Fig. 8). VLPs and EVs are very similar in composition, and it has been postulated that they use similar pathways for vesicle trafficking (58, 59). So, unlike S, ΔS was efficiently incorporated into VLPs or EVs like for example tetraspanins or endosomal markers that are equally found in both particle types (58, 59).

EVs released from 293GP-ΔS contain less than 10% of the total ΔS protein, and they would not need to be removed from vaccine preparations as they could be as good immunogens as VLPs. It was even reported that EVs containing the S protein of SARS-CoV could induce high levels of Nabs (60).

Titers of recombinant GFP retroviruses released from stable producers were at least a 1000-fold higher with ΔS versus S despite a 15-fold difference in the amount of the two proteins incorporated at the surface of VLPs (Fig. 5A and Fig 7B). As we did not find major differences in fusogenicity between S and ΔS in a syncytia formation assay (Fig. 6), our results suggest that recombinant viruses become fully infectious when a certain threshold of S protein is incorporated at their surface.

Recombinant GFP or luciferase pseudotyped retroviruses are commonly used to measure the activity of Nabs present in serum of infected or vaccinated people (55–57). These reagents are convenient, as unlike SARS-CoV-2 they can be manipulated in a BSL-2 laboratory. The robust production system with the 293GP-ΔS cell line could be highly valuable to evaluate the presence of Nabs in large cohorts.

Mass production will be a major challenge with all types of SARS-CoV-2 vaccine that are being developed as the entire worldwide population will have to be vaccinated. Based on the results of a nanoparticle vaccine containing S, whose 5 and 25 *μ*g doses triggered a high level of Nabs in people (28), we assume that a vaccine derived from the VLP platform described in this study could be efficient with similar or lower amounts of S per dose. The yield of VLPs produced from the 293GP-ΔS cells could be increased if a high producer clone is selected instead of a bulk population, and if cells are cultured in bioreactors in fed-batch or perfusion modes. The average titer of gene therapy vectors produced with a derivative of the 293GP cell line was increased by 5.6-fold in bioreactor versus a 10-layer cell factory, and the total vector yield was increased by 13.1-fold (61). Mutations of the furin cleavage site located between S1 and S2 and the D614G variant that is now more prevalent in the infected population could increase the amount of S incorporated into VLPs (57, 62).

A very concise review that compared the first results of different COVID-19 vaccines concluded that the most immunogenic ones were made with recombinant proteins (25). These results emphasize the importance of the platform developed in this study because VLPs present the antigen in a protein format that seems more potent for vaccination than the protein alone. Indeed, MLV VLPs displaying the human cytomegalovirus glycoprotein B antigen could trigger 10-times more Nabs in mice than the protein alone using the same amount of antigen (63). Finally, VLP-S could be used as a boost for other types of vaccine like measle virus- and adenovirus-based recombinant vectors. These combinations were highly potent for triggering Nabs against hepatitis C proteins in mice and macaques (64).

In conclusion, we have developed and characterized a new MLV VLP platform that can efficiently incorporate the S protein from SARS-CoV-2, and that has the potential to produce a pan-coronavirus vaccine. The next logical step is to validate this vaccine in experimental animals and in humans thereafter.

## MATERIALS AND METHODS

### Plasmids

The expression plasmid pMD2ACE2iPuro^r^ containing the human angiotensin-converting enzyme (ACE2) cDNA used to generate ACE2 positive cells was constructed as follows: the ACE2 *PmeI* cDNA fragment obtained from the plasmid hACE2 (Addgene; #1786) was cloned in pMD2iPuro^r^ opened in EcoRV.

The SARS-CoV-2 *S* gene from the Wuhan-Hu-1 isolate (GenBank: MN908947.3) was codon optimized (Genscript, Township, NJ) and cloned in pMD2iPuro^r^ in *EcoRI/XhoI*. A shorter version with a 19-codon deletion in C-terminal (ΔS) was also constructed in a similar way.

The pMD2.GalviPuro^r^ and pMD2.G plasmids that encode the Galv and VSV-G envelopes, and the retroviral vector plasmid containing the *GFP* gene under the control of the 5’ long terminal repeat sequence have been described elsewhere (65).

### Cell Lines

293GP, 293 cells (ATCC, CRL-11268), and their derivatives expressing the ACE2 receptor (293-ACE2), S (293GP-S and 293-S), DeltaS (293GP-ΔS and 293-ΔS) and the Galv envelope (293GP-Galv) were cultured with Dulbecco’s modified Eagle’s medium (DMEM; Wisent, Canada) supplemented with 10% fetal calf serum (Life Technologies, Grand island, NY) and antibiotics (Wisent). Bulk populations of 293-ACE2, 293-S, 293-ΔS, 293GP-S, 293GP-ΔS and 293GP-Galv were established by transfection using the calcium phosphate procedure. Briefly, subconfluent 293 or 293GP cells plated in 10-cm dishes were transfected with 20 μg of the pMD2 plasmids expressing ACE2, S, ΔS or Galv. Two days later, cells were selected in puromycin for a period of 10 days (0.5 μg/ml). Bulk populations of 293GP-S/GFP, 293GP-ΔS/GFP and 293GP-Galv/GFP were generated by infections of the parental cells with a GFP vector pseudotyped with VSV-G produced in transient transfection. The 3 derived cell lines were at least 86% GFP positive (Fig. 4B).

### Virus Productions and Infections

The production of GFP recombinant retroviruses was generated by transient transfection of 293 cells. One day prior transfection, 3 × 10^6^ cells were plated in 60-mm dishes. 293 cells were transfected for 4-hours by the calcium phosphate procedure with 1 μg of envelope expression plasmids (pMD2.G, pMD2.GalviPuro^r^, pMD2.SiPuro^r^ or pMD2.ΔSiPuro^r^), 4 μg of Gag-Pol expression plasmid (pMD2GPiZeo^r^) and 5 μg of RetroVec plasmid. Two days later, 2.5 ml of viral supernatants were harvested and frozen at −80°C. Recombinant viruses from stable 293GP-S/GFP, 293GP-ΔS/GFP and 293GP-Galv/GFP cells were also produced similarly in 60-mm dishes.

Transduction efficiency of GFP vectors was determined by scoring fluorescent-positive target cells. 293-ACE2 cells were inoculated at a density of 2 × 10^5^ cells per well in 24-well plates. The next day, the medium from each well was replaced with different volume of viral supernatants containing 8 μg/ml polybrene. Two days later, cells were trypsinized and analyzed by flow cytometry. Vector titers were calculated using the following formula (N × P) × 2/(V × D). N= Cell number on the day of infection; P= percentage of fluorescent-positive cells determined by flow cytometry; V is the viral volume applied; and D is the virus dilution factor. Titers were calculated when the percentage of fluorescent-positive cells was comprised between 2 to 20%. Alternatively, GFP positive cells were assessed under a fluorescent microscope. The 3 × 3 mosaic images of GFP and transmitted light were acquired with a Nikon TI-E inverted microscope with a PlanApo VC 20x 0.75 NA objective using a Hamamatsu Orca-ER CCD camera. Acquisition and stitching were performed with the Nikon NIS Elements 5.02 software program. The fluorescence intensity of infected cells displayed in figure 5A were scanned using the Fiji software to evaluate the difference in viral titers (66).

### Syncytia Formation Assay

293-ACE2 cells were mixed with 293, 293GP-S and 293GP-ΔS at a 9/1 ratio and plated at 4 × 10^5^ cells/well in a 24-well plate. Fusion activity was analyzed 24 h later by phase contrast under the same microscope used for measuring the transduction efficiency.

### Protein Analysis

The presence of S at the surface of 293 cells was assessed in transient transfections. Subconfluent cells in 6-well plates were transfected for 4 hours with 5 *μ*g of S or ΔS plasmids by the calcium phosphate procedure, and 24 hours later, the media was replaced with serum-free media (SFM) BalanCD HEK293 (Fujifilm Irvine Scientific, Santa Ana, CA). The next day, cells were detached without trypsin by gently pipetting up and down the medium on top of the cells. A human chimeric anti-S1 antibody (Genscript; 1:200 dilution) followed by an Alexa647-conjugated goat anti-human IgG (Jackson Laboratories; 1:400) were successively incubated with cells for labelling. The fixable viability stain 450 (BD Biosciences, San Jose, CA, USA) was used to exclude dead cells. The presence of S was then analyzed by flow cytometry with a BD FACSAria II (BD Biosciences). Cells transfected with a Galv expression plasmid were used as control. The presence of stably expressed S at the cell surface of 293GP-S and 293GP-ΔS was similarly analyzed by flow cytometry.

The presence of ACE2 at the surface of 293-ACE2 cells was also checked by FACS. Detached cells were labelled with a mouse anti-ACE2 antibody (R&D Systems, Minneapolis, MN1/200) followed by an Alexa488 goat anti-mouse (1:1,000; Invitrogen, Carlsbad, CA).

The presence of S released in the supernatant of transiently transfected 293GP cells was analyzed by Western blot. Subconfluent cells plated in 60 mm were transfected for 4 h with 5 *μ*g of envelope expression plasmids and 5 *μ*g of the GFP retroviral plasmid. One day later, the media was replaced with 2.5 ml of SFM that was then harvested the following day. Supernatants were concentrated 10-fold with a 30 kDa Amicon centrifugal unit (Millipore Sigma, Oakville, Canada) and were stored at −80°C until use. The GFP fluorescence evaluated under a microscope at the time of harvest was very similar among the different transfected plates.

Supernatants from confluent 293GP-S, 293GP-ΔS, 293-S and 293-ΔS cells were also harvested and concentrated from 60-mm dishes.

Cell pellets of 1 × 10^6^ cells were resuspended in 100 *μ*l RIPA lysis buffer containing a protease inhibitor cocktail (Roche). Samples were centrifuged for 5 min to remove cell debris and stored at −20°C until use for Western blot analysis.

Samples of 20 *μ*l were incubated 5 min at 95°C in loading buffer containing 1% SDS and 2.5% β-mercaptoethanol, and run on a 10% SDS-polyacrylamide gel (4% stacking), followed by transfer onto nitrocellulose membranes (GE Healthcare). Immunoblotting was performed with a rabbit polyclonal antibody anti-S2 (1:400 dilution, SinoBiological, Beijing, China) and a rat monoclonal antibody anti-MLV p30 produced from the hybridoma R187 (1:2,000 dilution; American Type Culture Collection, Manassas, VA). Blots were then incubated with secondary antibodies IRDylight680 goat anti-rat IgG (1:10,000; Invitrogen) and IRDye 800CW anti-rabbit IgG (1:10,000; Li-Cor Biosciences, Lincoln, NE), and analyzed with the Odyssey Infrared Imaging System (Li-Cor Biosciences). Serial dilutions of known amounts of C-terminally Fc-tagged S2 (BioVendor, Brno, Czech Republic) were used for quantification.

### Velocity Gradient

Thirty ml of supernatant from confluent 150-mm dishes of 293GP-ΔS and 293-ΔS cells were harvested and filtered through a 0.45 *μ* membrane, and concentrated by ultracentrifugation for 90 min at 100,000 × g in a AH629 rotor. Pellets containing virions and EVs were resuspended in 1 ml PBS containing a protease cocktail inhibitor (Roche) during 2 h at 4°C. The resuspended vesicles were layered onto a 6-18% Optiprep™ 11-step discontinuous velocity gradient (Stemcell Technologies, Vancouver, Canada), and centrifuged for 90 min at 176,000 × g in a SW40Ti rotor as previously described (48, 49). Fractions of approximately 800 *μ*l were collected from the bottom after puncturing the wall of the centrifuge tube with a gauge needle, and 20 *μ*l of each sample were analyzed by Western blot.

## ACKNOWLEDGMENTS

The authors thank Carl Saint-Pierre for his technical assistance in microscopy.

This work was supported by BioVec Pharma. KG, POdeCL and MC are funders and shareholders of BioVec Pharma. MC is an author of a patent application covering the VLP platform presented in this study.

